# No Correlation Between Pollen Fertility and Viability: Differential Measures of Male Gametophytic Fitness in *Cannabis sativa* L.

**DOI:** 10.1101/2021.11.04.467333

**Authors:** Sydney B. Wizenberg, Michelle Dang, Lesley G. Campbell

## Abstract

Pollen grains are male gametophytes, an ephemeral haploid generation of plants, commonly engaging in competition for a limited supply of ovules. Since differential male fertility may influence the direction and pace of population evolution, the relative fitness of pollen is regularly estimated as either pollen viability, the proportion of pollen containing intact cytoplasm’s and regenerative nuclei, or pollen fertility, the frequency of pollen germinating under standardized conditions. Here, we estimated the relative fitness of pollen in a dioecious, wind-pollinated model system, *Cannabis sativa*, by characterizing pollen fertility and viability from multiple sires. Pollen fertility quickly declined within two weeks of anther dehiscence, and pollen stored under freezer conditions did not germinate regardless of storage time. In contrast, pollen viability declined slowly and persisted longer than the lifetime of a sporophyte plant under both room temperature and freezer conditions. Pollen samples that underwent both fertility and viability analysis displayed no significant correlation, implying researchers cannot predict pollen fertility from pollen viability, nor infer male gametophytic fitness from a single measure. Our work demonstrates two approaches to measure proxies of male fitness in *C. sativa*, and identifies new questions around what are valuable estimates of male fitness in plants.

## Introduction

Pollen grains are male gametophytes, representing an ephemeral, haploid generation in the plant life cycle (Hafidh et al., 2016; Maheshwari, 1949; Walbot and Evans, 2003). Pollen compete for opportunities to fertilize ovules and thus, influence differential reproductive success, a key condition for microevolution (Beaudry et al., 2020; Johnson and Shaw, 2016; Ottaviano et al., 1988; Ottaviano and Mulcahy, 1986). Post-meiotic gene expression and the abundance of pollen produced by many gymnosperms and angiosperms sets the stage for intense competition among pollen-derived sperm cells for the often limited supply of ovules, wherein sporophyte reproductive success relies on not only successful dispersal of pollen, but also successful pollination of stigmas and subsequent pollen germination and ovule fertilization (Ottaviano et al., 1988; Ottaviano and Mulcahy, 1986). To better understand and explore the competition occurring at the stage of pollen production and release, we must characterize and estimate their relative fitness, including: survival and fertility (Ottaviano et al., 1982). While there is a substantial body of literature estimating the relative fitness of pollen (Choi et al., 2014; Hauser and Siegismund, 2000; Hudson and Stewart, 2004; Luria et al., 2019; McDowell et al., 2015), the terminology used to describe these characteristics is inconsistent. Though some authors correctly differentiate between fertility and viability (Barrow, 1983; Dafni and Firmage, 2000; Rajasekharan and Ganeshan, 1994; Trognitz, 1991), others use the terms interchangeably as synonyms, reflecting broad misconceptions of these terms (Qureshi et al., 2009; Shukla et al., 2020; Tuinstra and Wedel, 2000). Definitively, viability measures cytoplasmic degradation of the regenerative cell to quantify the proportion of pollen from a sire containing intact regenerative nuclei (Alexander, 1969; Atlagić et al., 2012; Barrow, 1983; De Souza et al., 2003; Firmage and Dafni, 2000; Heslop-Harrison and Heslop-Harrison, 1970; Impe et al., 2020; Khatun and Flowers, 1995; Pinillos and Cuevas, 2008; Rao et al., 1992; Rich, 2009; Rodriguez-Riano and Dafni, 2000; Shivanna and Heslop-Harrison, 1981). Viability measures how many pollen grains are capable of engaging in reproduction under any condition, as a degraded cytoplasm indicates senescence and death of the gamete, as opposed to measuring how many pollen grains may germinate under a particular set of conditions. Comparatively, fertility is quantified through measurement of the proportion of pollen germinating under standardized conditions (Adaniya and Shirai, 2001; Conner, 2011; Heslop-Harrison, 1987; Janssen and Hermsen, 1976; Jayaprakash, 2018; Říhová et al., 1996; Sharma et al., 1990; Soares et al., 2008; Yates and Sparks, 1990). Successful germination of a pollen grain requires rehydration of the apertures and subsequent protrusion of the pollen tube towards the ovule, following which the pollen tube transports the male gamete to the female ovule to produce a zygote (Heslop-Harrison, 1987; Taylor and Hepler, 1997). Both pollen grain viability and fertility are used as measures of the proportion of pollen capable of engaging in reproduction under standardized conditions, and quantification of these measures could provide insight into why, or how, some male plants possess reproductive advantages.

Viability can be tested using many methods (Atlagić et al., 2012; Pinillos and Cuevas, 2008; Rodriguez-Riano and Dafni, 2000; Trognitz, 1991), but one common means of differential staining is the Alexander stain. Originally published in 1969, the Alexander stain uses acid fuchsin and malachite green to test if the cytoplasm containing the regenerative nucleus is intact (Alexander, 1969). Malachite green stains the exine and intine cell walls blue, while acid fuchsin is absorbed by the cytoplasm, resulting in a pink stain (Alexander, 1969). Viable and non-viable pollen are differentiated by the resulting coloration, wherein pollen staining pink within the vegetative or regenerative cell did not contain an in-tact regenerative nucleus, and were therefore incapable of fertilizing an ovule regardless of the external germination conditions (Alexander, 1969). Originally containing chloral hydrate, mercuric chloride, and phenol, a simplified version of the Alexander stain was developed and tested in 2010, allowing wider applications of this method due to the removal of some toxic and difficult to acquire chemical components (Peterson et al., 2010). This simplified method was successful at differentiating between viable and non-viable pollen in numerous test species (*Gingko biloba, Pinus resinosa, Acer rubrum, Arabidopsis thaliana, Betula populifolia, Fragaria versca, Lonicera tatarica, Oryza sativa, Prunus padus, Rhododendron mucronulatum*) and shows promise in its ability to act as one standardized, interspecific method of differential pollen viability staining (Peterson et al., 2010). Measuring pollen fertility through *in vitro* germination is more challenging to standardize, as germination medias are typically developed for individual species, based on their distinct biochemical germination signals (Jayaprakash, 2018; Patel and Mankad, 2014). Some components, such as water, sucrose, boric acid, and polyethylene glycol, are used frequently in medias developed for different species, while other growth inducing additives such as calcium chloride, potassium chloride, potassium nitrate, and magnesium sulphate, can vary significantly in their quantity (Acar et al., 2010; Ateyyeh, 2005; Díaz and Garay, 2008; Fragallah et al., 2019; Imani et al., 2011; Jayaprakash, 2018; Jayaprakash and Sarla, 2001; Mercado et al., 1994; Mortazavi et al., 2010; Potts and Marsden-Smedley, 1989). All germination medias must contain a combination of carbohydrate sources in addition to growth inducing additives to mimic biochemical indicators of stigma proximity and induce rehydration and subsequent pollen tube growth (Heslop-Harrison, 1987; Jayaprakash, 2018). Because of variation in germination medias composition among species, standardization is difficult, and prevents relative comparisons of fertility between species relying on diverse germination medias.

*Cannabis sativa* L. is a dioecious crop frequently cultivated for its cannabinoids, fibre, and seeds (Amaducci et al., 2008; Bertoli et al., 2010; Carus and Sarmento, 2016; Grégoire et al., 2020; Small, 2015; van derv Werf et al., 1996; Zuardi, 2006). This species is anemophilous or wind-pollinated, and its exine morphology reflects this dispersal strategy, meaning its pollen grains are not ornamented and thus well suited to rapid movement coinciding with any changes in air flow (Halbritter et al., 2007; Small and Antle, 2003). Indoor industrial facilities producing cannabinoids from *C. sativa* often produce sinsemilla, unpollinated floral biomass, because pollination of female plants causes a reduction in cannabinoid content and the length of trichome dense stigmas (Ohlsson et al., 1971; Valle et al., 1968). Regulations relating to the production of *C. sativa* in countries such as Canada often include specific stipulations requiring that floral biomass is seedless (sinsemilla), therefor growing female plants in strict isolation is both favorable and legally required in some regions (Government of Canada, 2019).

In addition to answering basic scientific questions about male gametophytes and their life cycle, investigating the behaviour and related characteristics of pollen in this economically valuable species could improve our ability to control pollination risk within *C. sativa* growing facilities. Zottini *et al.* investigated the effect of gamma ray irradiation on pollen viability and fertility, using loaded fluorescein diacetate to differentiate between viable and non-viable grains under a microscope, and 5 different germination medias (Zottini et al., 1997). They found gamma ray irradiation did not affect measures of viability, but drastically reduced rates of *in vitro* germination, and noted estimates of fertility and viability differed from one another (Zottini et al., 1997). Rana and Choudhary (2010) tested three different measures of viability (Alexander’s stain, triphenyl tetrazolium chloride, and flurochromatic reaction) and documented a substantial decline in viability within three days of anther dehiscence (Rana and Choudhary, 2010). More recently, Gaudet *et al.* (2020) developed a media for *in vitro* germination and investigated long-term cryopreservation of *C. sativa* pollen for use in breeding programs. Pollen collected at diverse developmental stages also differed in its germination capabilities; some samples maintained fertility past three weeks, while others declined rapidly in the first two weeks of storage (Gaudet et al., 2020). Pollen stored in wheat flour and liquid nitrogen maintained its ability to germinate *in vitro* for up to four months, though the germination rates remained low, even for fresh pollen (Gaudet et al., 2020). Noting the inconsistencies in methodologies and conclusions among previous publications on this topic, and building on our previous work (Wizenberg et al., 2020), we set out to investigate *C. sativa* pollen viability and fertility, under the broader goal of developing a framework for measuring male gametophytic fitness. Accordingly, our research asks:

1. How long is *C. sativa* pollen viable under standard and freezer conditions?
2. How long is *C. sativa* pollen fertile under standard and freezer conditions?
3. Can measures of *C. sativa* pollen viability predict pollen fertility?

## Results

Pollen fertility and viability were generally high on the first day they were measured and declined with time (see below). Samples tested directly after collection had an average viability of 93.35 % (± 3.74%) and an average fertility of 38.69% (± 3.15%). Pollen stored under room temperature conditions (20 °C, 43% RH) showed a consistent decline in viability over the 12-week monitoring program, and the associated linear model predicted pollen samples under these conditions may maintain viability for up to 39 weeks, approximately nine months post anther dehiscence. The linear model fitted to predict pollen viability when stored under room temperature conditions was showed a strong model fit (Adj. R^2^ = 0.89, RSE = 3.12, df = 33, F = 277, p < 0.001; Fig. 1a). Pollen stored under freezer conditions (−4 °C and ~100% RH) maintained high viability even after 96 weeks of storage, and the associated linear model predicted pollen stored under freezer conditions may maintain viability up to 261.5 weeks, approximately five years after anther dehiscence. The linear model fitted to predict pollen viability when stored in a freezer also showed a strong model fit (Adj. R^2^ = 0.89, RSE = 3.79, df = 23, F = 214.8, p < 0.001; Fig. 1b).

**Figure 1:**
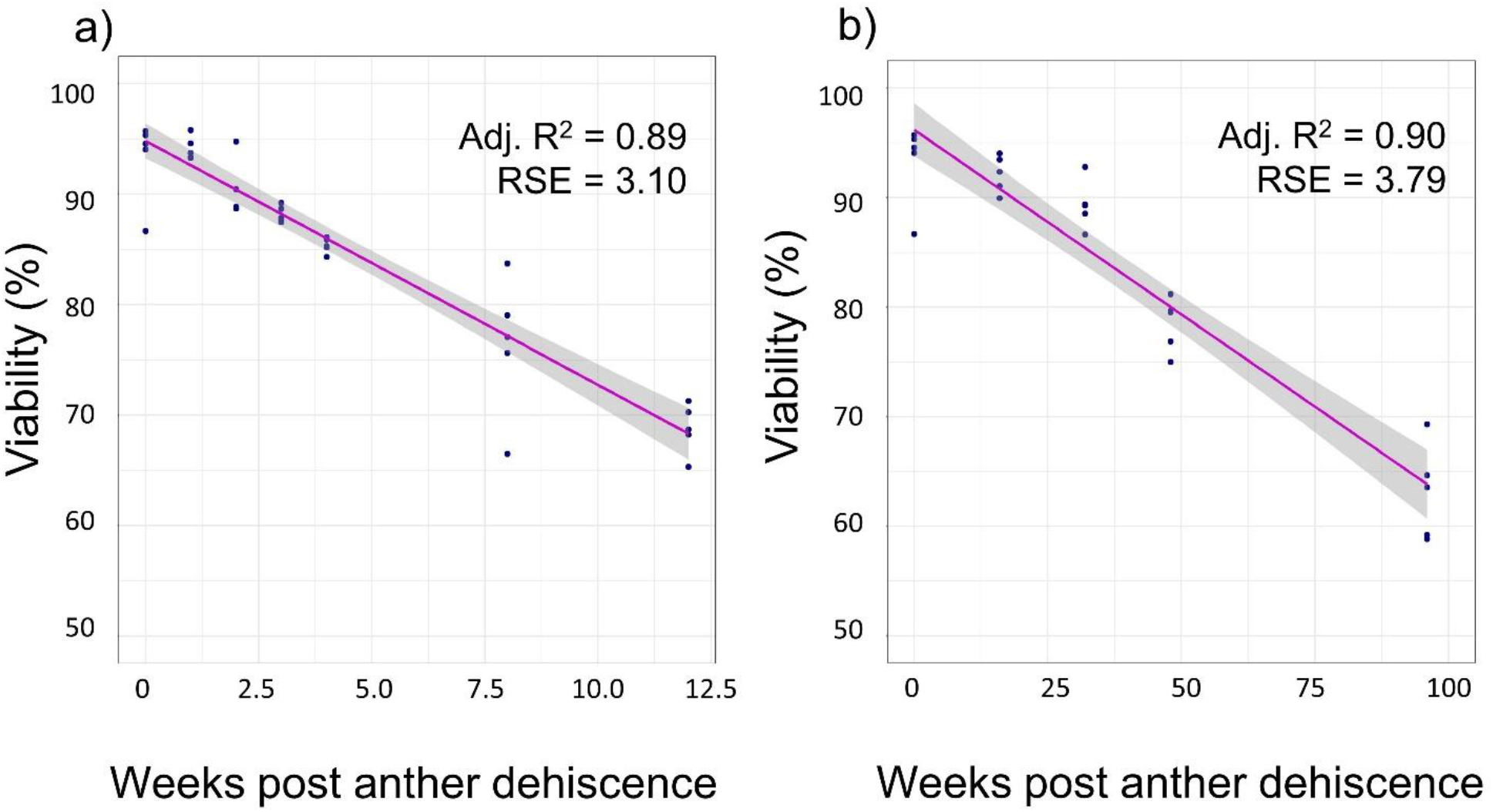
Pollen grain viability under two experimental conditions. The linear relationship between weeks post anther dehiscence and viability of pollen stored under (a) room temperature conditions (22 ± 0.95 C, ~43% r.h.) and (b) freezer temperature conditions (−4 °C, ~100% r.h.).

Pollen stored under room temperature conditions (20 °C, 43% RH) quickly declined in fertility, reaching 0% germination two weeks after anther dehiscence. The linear model fitted to predict pollen fertility when stored under room temperature conditions showed a moderate model fit (Adj. R^2^ = 0.87, RSE = 6.34, df = 13, F = 92.67, p < 0.001; Fig. 2). Pollen stored in a freezer (−4 °C and ~100% RH) showed no germination regardless of storage time, after testing *in vitro* germination at both one and two weeks and seeing no pollen tube growth we discontinued collecting data on fertility of frozen pollen.

**Figure 2:**
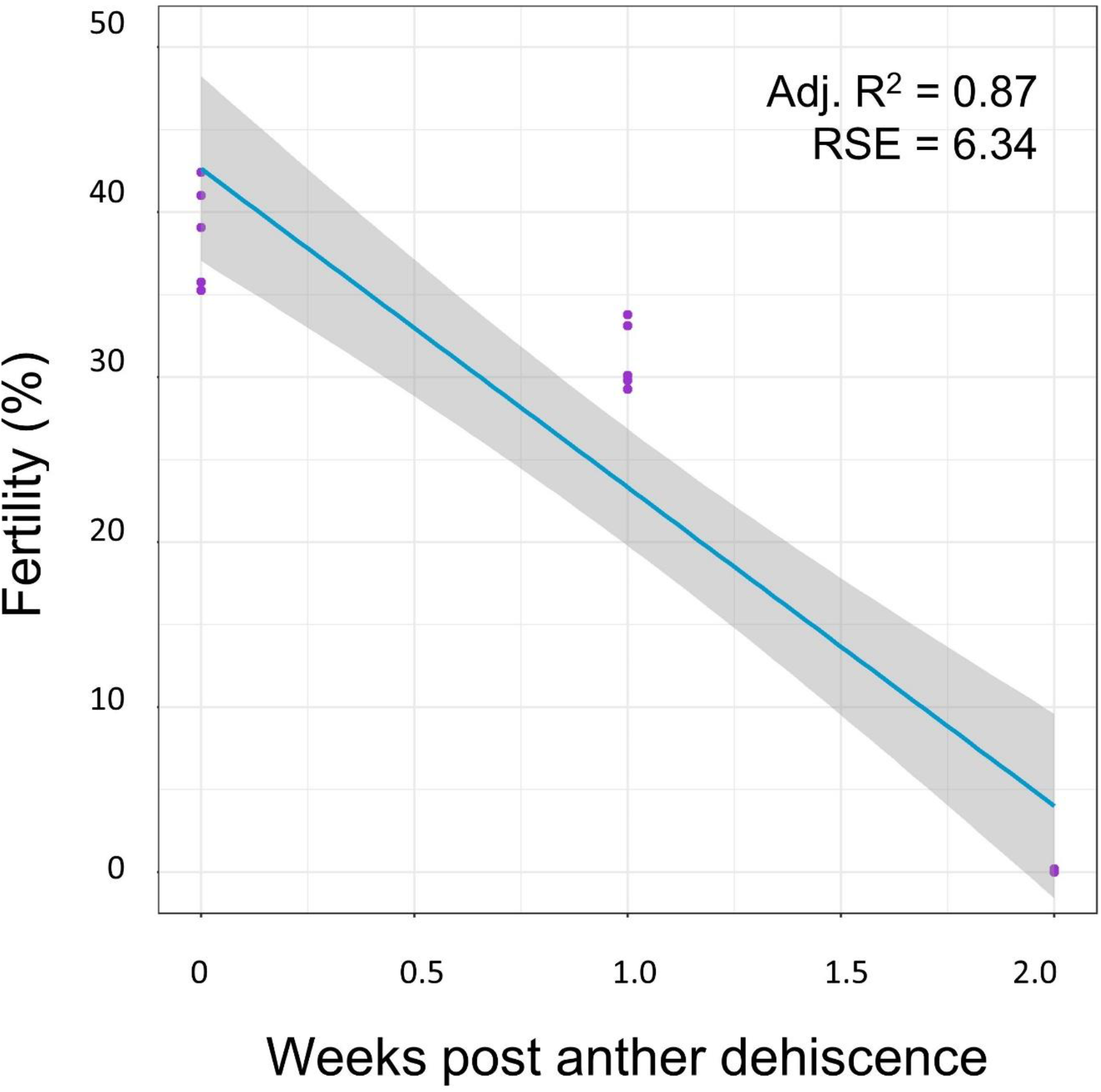
Pollen grain fertility under room temperature conditions. The linear relationship between weeks post anther dehiscence and the fertility of pools of pollen grains stored under room temperature conditions (22 ± 0.95 C, ~43% r.h.)

Pollen fertility did not significantly predict pollen viability using a linear model, when both traits were measured for a single sample (Adj. R^2^ = −0.06, RSE = 17.93, df = 13, F = 0.23, p = 0.64). Moreover, paired measures of pollen fertility and viability showed no detectable association using non-parametric approaches to analysis (Wilcoxon signed rank test: v = 120, p < 0.001), further demonstrating viability and fertility are independent measures of male gametophytic fitness and do not express any dependent linear relationship (Fig. 3).

**Figure 3:**
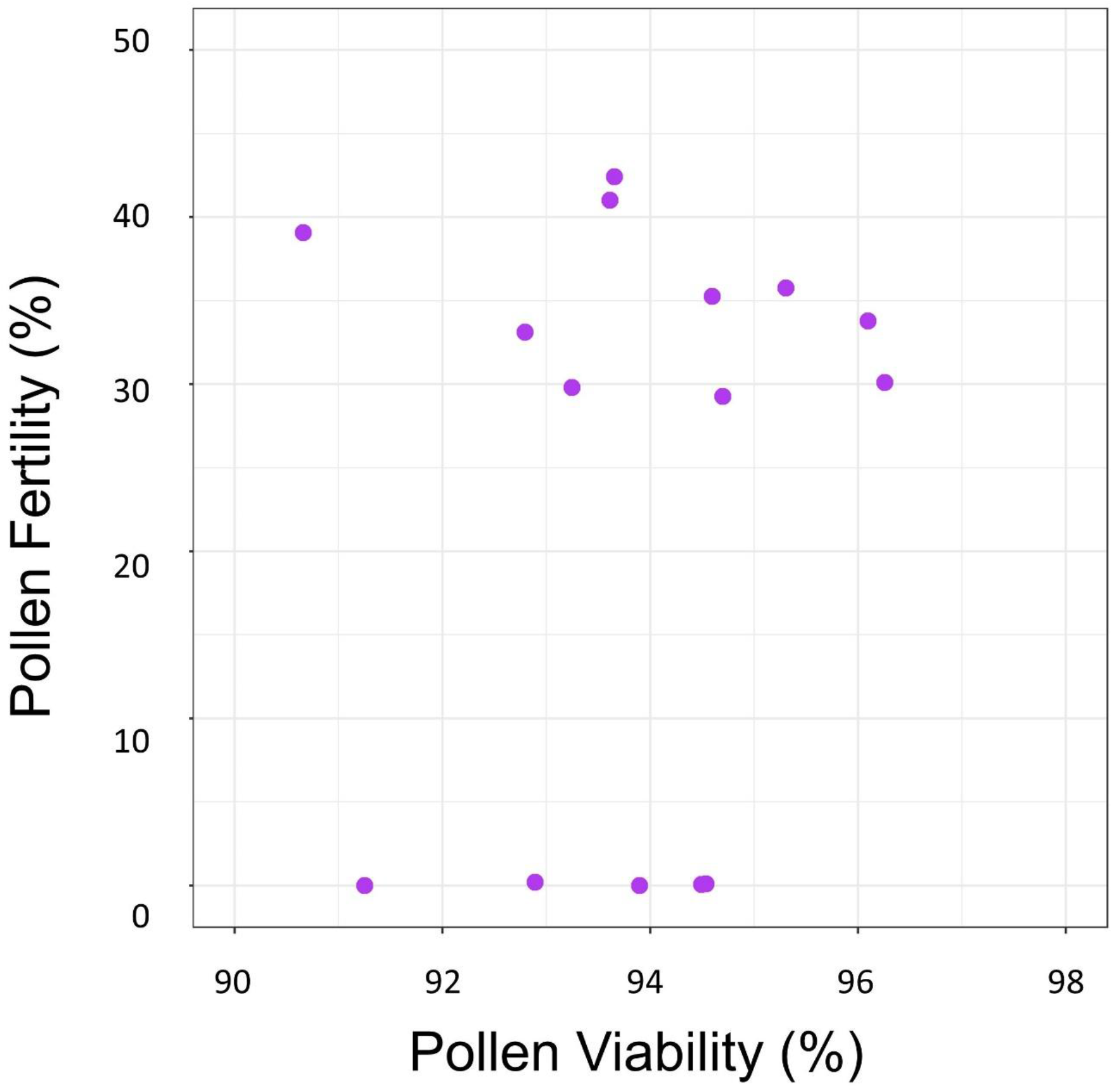
Pollen grain viability does not predict fertility. The relationship between fertility and viability of pollen grains from a single sample. The associated linear model has insufficient model fit (Adj. R^2^ = −0.06, RSE = 17.93); Wilcoxon signed rank analysis determined that the two characteristics were independent, showing no detectable relationship (v = 120, p < 0.001).

## Discussion

Our work has demonstrated fertility and viability, two differential measures of male gametophytic fitness, are independent of each other and show no correlation in *C. sativa*. This coincides with previous *C. sativa* research by Zottini *et al*., wherein they also found no relationship between these characteristics (Zottini et al., 1997). This may be due, in part, to contrasting sensitivity to storage conditions and the external environment; *in vitro* germination requires rehydration and pollen tube growth, a complex biochemical and physiological process (J Heslop-Harrison, 1987). Comparatively, measures of viability rely on pollen senescence, a simpler process that may be less sensitive to storage conditions and the external environment (Gahan, 1981). Pollen fertility appeared to be more sensitive to environmental degradation than viability, as it quickly declined within two weeks of anther dehiscence, and pollen stored in the freezer (−4 °C and ~100% RH) did not germinate regardless of storage time, potentially as a result of the high humidity preventing sufficient dehydration. Interestingly, fresh pollen incubated in a germination media directly from dehiscent anthers did not exceed 42.4% germination, with an average of 38.7% at this time point, similar to the results obtained by Gaudet *et al.,* who developed the media recipe and found fresh pollen germinated at rates varying between 30-50% depending on phenological behaviour (Gaudet et al., 2020). This could imply fertility is generally low in *C. sativa*, or the germination media could be further optimized to reach germination rates exceeding 50%. Previous research investigating pollen fertility in other species has demonstrated *in vitro* germination rates can exceed 80% through optimization of the content of sucrose, polyethylene glycol, and other growth inducing additives (Jayaprakash and Sarla, 2001), while others have shown relative humidity can strongly influence maximum germination rates under standardized conditions (Burke et al., 2004). More recently, optimization of *in vitro* germination in another crop (*Pheonix dactylifera*) has also failed to exceed a threshold of 60%, implying *C. sativa* is not the only species struggling to germinate in liquid media (de Oliveira et al., 2021). Bearing this in mind, quantitative measures of fertility in a germination media may not reflect real life fertility rates, but rather may only be used as a relative indicator of reproductive performance.

Viability under room temperature conditions reached a maximum of 95.8%, and averaged 93.3% for freshly dehiscent pollen. Thus, most fresh pollen produced by male *Cannabis* plants are capable of engaging in reproduction under optimal conditions, and very few pollen grains do not contain an intact regenerative nucleus following maturation of the pollen sacs. Pollen viability under room temperature conditions averaged 68.8% 12-weeks after anther dehiscence, exceeding the approximate length of the *Cannabis* life cycle (Small et al., 2003). Thus, pollen senescence is a process that may be manipulated to preserve pollen samples for extended periods of time. This is verified by the high viability of pollen grains stored in the freezer for 96 weeks post anther dehiscence, which averaged 63.1% viability, with a maximum of 69.3% and a minimum of 58.8%. The linear regressions used to model the relationship between storage time and viability predicted that pollen would lose all viability at 38.3 weeks under room temperature conditions (Adj. R^2^ = 0.8903), and 261.5 weeks under freezer conditions (Adj. R^2^ = 0.8991), suggesting that long-term storage of pollen samples for genotyping is feasible. Furthermore, any pollen that escapes into growth rooms may remain viable, long after the crop has been harvested. However, the steady decline in fertility that occurred under both experimental conditions could prevent use of stored pollen samples for breeding programs, though it may be possible to extend the decline in fertility if storage conditions and the germination media were further optimized.

Investigation of the relationship between pollen viability and fertility in other species has yielded results largely similar to our conclusions. Trognitz (1991) explored correlation between viability assays and proxies of fertility in potatoes, finding that some methods of viability assessment predicted fertility more strongly than others (e.g., enzyme activity assays were a good predictor of pollen fertility but pollen staining by carmino acetic acid was not). Other research in cotton found that both pollen staining and germination were poor indicators of over-all fertility for this species, but under multiple storage conditions, pollen that stained as viable also germinated *in situ* and in an agar media (Barrow, 1983). Barnabas and Rajki (1981) tested the utility of year old *Maize* pollen, finding it to maintain 50% viability and 30% fertility, indicating that fertility declines much more rapidly than viability in this species as well. In bananas, measures of fertility and viability differed in multiple genotypes and under two different germination media compositions, though these authors did not directly test any related hypotheses (Soares et al., 2008). More recently, Chhun et al. (2007) demonstrated that gibberellin regulates both characteristics, but in different ways, implying that these attributes are interrelated yet distinct. Broadly, measures of fertility and viability differ across species and between characterization methods, through some work has indicated that correlation exists (Trognitz, 1991) depending on the individual procedures and techniques.

Through development and testing of these methods for characterizing male gametophytic fitness in *C. sativa* we have shown that the pollen of this species remain viable for extended periods of time but do not maintain fertility under these conditions beyond two weeks after anther dehiscence. Though these results are promising, some limitations of our work prevent larger inferences relating to male gametophytic fitness as a whole. Quantitative fertility, measured through *in vitro* germination, may not reflect real world germination rates, and can therefore only be used for relative comparisons between sires. The long-term viability documented in this experiment does not coincide with previous research on this species (Rana and Choudhary, 2010), potentially as a result of differences in methodology or genotypes used. Additionally, the use of early flowering males limits our ability to investigate if phenological behaviour influences fertility, as documented by Gaudet et al. (Gaudet et al., 2020). Future work on this topic should include multiple genotypes and pollen collected at different developmental checkpoints to investigate how these factors influence measures of male gametophytic fitness.

## Methods

Over two years (2019-2021), we developed and tested a framework for measuring the relative fitness of pollen characterized by two traits: pollen grain viability and pollen grain fertility. We tested the Alexander stain to measure pollen viability (Alexander, 1969; Peterson et al., 2010) and Gaudet’s media to measure *in vitro* germination (Gaudet et al., 2020) on the pollen of *C. sativa* to establish baseline relative fitness values and measure pollen survival through time. Additionally, we tested if pollen grain viability and fertility were correlated when samples were collected from the same pollen source.

### Plant genotype and cultivation methods

To test pollen viability and fertility hypotheses, we grew CFX-2 (Hemp Genetics International, Saskatoon, Saskatchewan, Canada), a hemp cultivar of *C. sativa* with an expected total tetrahydrocannabinol (THC) content of less than 0.01%. The project was divided into three distinct experiments, which started on May 29^th^, 2019; October 21^st^, 2019; and April 2^nd^, 2021, respectively. For each experiment, we germinated 20 seeds in a terracotta germination pot (ANVÄNDBAR Sprouter; IKEA, Delft, The Netherlands) for three days, watering once daily with 25 mL of filtered water (Milli-Q purification system #F7KA48180D, Millipore Canada Ltd., Etobicoke, Ont.). Following germination, we planted all 20 seedlings into SC-10 cone-tainers® (Stuewe and Sons Inc., Tangent, Oregon, USA) filled with 200 mL of moistened PRO-MIX mycorrhizae peat moss growing medium (Premier Tech, Riviere-du-Loup, Quebec, Canada). Seven days later we transplanted the seedlings into circular pots (15 cm. diameter × 11 cm. height) filled with approximately 1 L. of moistened PRO-MIX mycorrhizae peat moss growing medium. We then placed the seedlings under 24h lighting from fluorescent T8 bulbs (F32W, Canarm lighting and fans, Brockville, Ontario, Canada) for four weeks, following which we switched to a 12h lighting photoperiod regimen to induce flowering for the remainder of the experiment. We watered each plant twice weekly with 50 mL of filtered water and fertilized them once weekly with 250 mL of 0.4% Miracle-Gro® (10-10-10 NPK; Scotts Miracle-Gro, Marysville, Ohio, USA) diluted in filtered water.

### Pollen collection

Once floral development was initiated, we identified and tagged ten pollen-producing plants using floral morphology, i.e., the visible development of pollen-producing inflorescences at apical branching junctions. Hereafter, we refer to these plants as males, though we did not perform genetic testing to confirm their karyotype. Of the ten pollen producing plants, we selected the five that flowered early for inclusion in the experiment to minimize any variation in pollen characteristics as a result of phenological differences (Gaudet et al., 2020). We closely monitored the five selected male plants until the inflorescences began to swell, showing visible protrusion of mature pollen sacs, indicating that anther dehiscence would occur. On the first day of anther dehiscence, we hand collected pollen (Wizenberg et al., 2020) in 1.7 mL centrifuge tubes (LIFEGENE graduated micro-centrifuge tubes, Modiin, Israel). Pollen samples in centrifuge tubes were left unsealed to dry for 1h before sealing, following which we stored them under one of two experimental conditions; samples were labelled and stored at either ‘room temperature’ conditions of 22 °C ± 0.95 °C and ~43% relative humidity, or ‘freezer’ conditions of −4 °C and ~100% relative humidity. For each male, we collected pollen twice on the initial day of anther dehiscence, with one sample being stored at ‘room temperature’ and the other being stored under ‘freezer’ conditions.

### Quantifying pollen viability

To test pollen viability, we prepared a batch of the modified Alexander stain (Peterson et al., 2010), and before the experiment, informally confirmed the presence of differential staining pollen containing functional cytoplasm (containing intact cytoplasm and a regenerative nucleus) or degraded cytoplasm (Fig. 4a). To evaluate the viability of each pollen specimen collected, we applied a small sample of pollen (ranging between 800 – 10,000 pollen grains), using a fresh cotton swab (Q-tips, Unilever, London, UK), to a 75 mm × 25 mm glass microscope slide. We pipetted 20 μL of the stain onto the pollen and heated the prepared slide 10 cm above a Bunsen burner for 5 s to set the stain. Once cooled, we applied a glass slide cover (25 mm × 25 mm) and the sample incubated at room temperature (22 ± 0.95 °C, 43% RH) for 24 hr. To estimate the proportion of pollen that stained viable as a percentage of the total sample, we counted the number of viable (stained pink) and aborted pollen (stained blue) using two hand-held tally counters (Uline, Pleasant Prairie, Wisconsin, USA) across vertical transects at 10x magnification (covering the entire length and width of the 2 mm.^2^ slide cover) under a compound microscope (Zeiss Primo Star Upright Light Microscope, Carl Zeiss Canada Ltd., Toronto, Ont.). Similarly to previous work (Wizenberg et al., 2020), we kept a tally of the number of burst pollen grains on each slide, separate from total pollen grains; burst pollen grains represented <1% of all sampled pollen grains. Any pollen forming clump were excluded from all counts because we could not differentiate between viable and aborted pollen. We measured pollen viability after initial anther dehiscence then weekly for the first four weeks of the experiment, and again after eight and twelve weeks for pollen stored at room temperature. We measured pollen viability after initial anther dehiscence then in 16-week intervals until 96 weeks for pollen stored at −4 °C.

**Figure 4:**
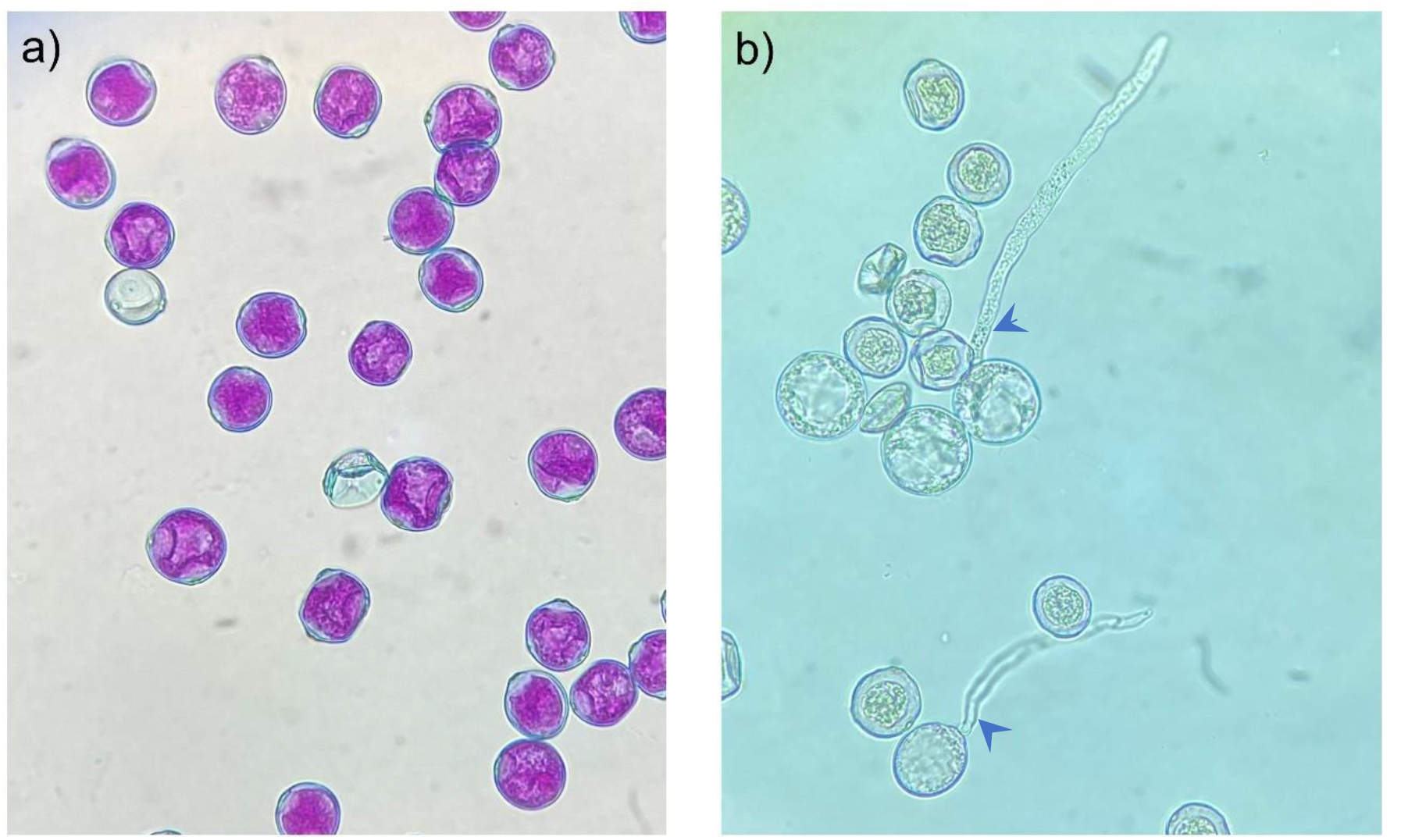
Differential staining and *in vitro* germination of *Cannabis sativa* pollen. (a) Differential staining of viable (pink cytoplasm with blue exines) and aborted pollen grains (blue exines), showing absorption of pink acid fuchsin in the functional cytoplasm of viable pollen grains. (b) *In vitro* germination of fertile and infertile pollen grains, showing protrusion of the pollen tube in fertile pollen grains (blue arrows).

### Quantifying pollen fertility

To test pollen fertility, we assembled a germination media recipe developed for *C. sativa* (Gaudet et al., 2020). Using a fresh cotton swab, we applied pollen (800 – 10,000 pollen grains) to a 75 mm × 25 mm glass microscope slide. We heated 10 mL of germination media in a 40 ml glass beaker placed on a hot plate at 70 °C for 10 min, at which point the media reached a temperature of 35 °C, then pipetted 50 μL of the heated media onto the pollen sample and covered it immediately with a glass slide cover (25 mm × 25 mm). We incubated the media at 28 °C (Fisher Scientific 6845 Isotemp Incubator 650D, Waltham, Massachusetts, USA) for 24 hr in a petri dish (8.5 cm × 8.5 cm × 1 cm) containing a filter paper (11 cm × 21 cm) soaked in 10 mL water to increase the relative humidity during incubation. After 24 hr, germination was visible (Fig. 4b) and we estimated the proportion of fertile pollen grains as a percentage of the total pollen sample using a compound microscope. We counted the number of germinated grains (rehydrated and containing a protruding pollen tube) and non-germinated grains across vertical transects at 10x magnification (covering the entire length and width of the 22 mm.^2^ slide cover). Again, we kept a tally of the number of burst pollen grains per slide, separate from the counted total; burst pollen grains represented <1% of all sampled pollen grains. Additionally, we excluded clumped pollen grains from all counts. We measured fertility of pollen stored at room and freezer conditions after initial anther dehiscence and then weekly until all samples showed no *in vitro* germination, at which point fertility was deemed to be zero.

### Statistical analyses

We conducted all analyses in R v.4.0.2, using the *stats* package (2019-04-06, R Core Team, 2019). We deemed all models to be sufficiently parametric using residual QQ plots. To determine if data collected conformed to typical patterns of pollen survival curves, we created three linear models where weeks post anther dehiscence predicted either viability (%) or fertility (%) using the ‘lm()’ function for separate data sets based on storage conditions. We gauged model strength through the adjusted R^2^, a goodness-of-fit statistics adjusted for the number of observations, and the residual standard error, both of which were reported using the ‘summary()’ function on each respective linear model. To measure the relationship between the paired estimates of pollen viability and fertility (from the parent pollen sample) we used a general linear model; the related summary statistics implied no dependent relationship (reported in the results), resulting in an insufficient model fit and weak predictive power. We transformed the data to investigate an inverse exponential relationship but this worsened the model fit and decreased the models predictive power. Finally, we investigated a potential non-parametric relationship between the two traits by performing the Wilcoxon signed-rank test, using the ‘wilcox.test()’ function contained in the *stats* package.

## Acknowledgments

The authors thank S. Sbrizzi and J. Muir-Guarnaccia for research assistance, and the department of Chemistry and Biology at Ryerson University for equipment. This work was supported by funds from the Natural Sciences and Engineering Research Council of Canada [DG #402305-2011 and DG #05780-2019 to LGC], as well as the department of Chemistry and Biology at Ryerson University. The funders had no role in study design, data collection and analysis, decision to publish, or preparation of the manuscript. Research was completed under the auspices of a federal hemp research license (#18-C0169-R-01) and a federal marijuana license (#LIC-U5GX543XM6).

## Author Contributions

SBW and LGC designed the experiment. SBW and MD prepared materials, developed methods, and collected data. SBW, LGC, and MD prepared the manuscript.

